# One-Dose Efficacy of Long-Acting Sorfequiline in a Mouse Model of Tuberculosis Preventive Therapy

**DOI:** 10.64898/2026.09.22.752266

**Authors:** Si-Yang Li, James J. Hobson, Nicole C. Ammerman, Henry Pertinez, Jo Sharp, Nader Fotouhi, Steve Rannard, Andrew Owen, Eric L. Nuermberger

## Abstract

**Rationale:** Tuberculosis preventive therapy (TPT) is essential for global tuberculosis control, yet estimated coverage remains well below global targets. Current World Health Organization (WHO)-recommended regimens require 1-9 months of treatment and are associated with suboptimal completion rates. Long-acting injectable (LAI) formulations offer the potential to improve TPT uptake and completion, with ideal products providing "one-and-done" options via a single administration. Sorfequiline, a second-generation diarylquinoline with superior potency over bedaquiline and lower QT prolongation risk, is a promising candidate for LAI TPT.

**Objectives:** To experimentally determine exposure-activity relationships for sorfequiline as TPT, estimate target plasma exposures, and characterize the pharmacokinetics (PK), tolerability, and efficacy of sorfequiline LAI formulations.

**Methods:** Oral sorfequiline dosing experiments established PK and pharmacodynamic relationships in a validated paucibacillary mouse model of TPT. Three candidate sorfequiline LAI formulations were evaluated for PK and tolerability in uninfected mice, followed by efficacy assessment compared to oral comparators (1HP, bedaquiline, sorfequiline).

**Measurement and Main Results:** Oral dosing experiments established a provisional target plasma C_min_ of 35 ng/mL for bactericidal effect. All tested LAI formulations achieved this target for at least 4 weeks post-dose. In efficacy studies, all sorfequiline LAI regimens (62.5-250 mg/kg) and sorfequiline oral regimens demonstrated significantly greater bactericidal activity than 1HP and oral bedaquiline. Among mice receiving ≥125 mg/kg of any sorfequiline LAI, 92% were culture-negative at 12 weeks.

**Conclusion:** Superior efficacy of a single-dose sorfequiline LAI injection compared to 1HP indicates the potential to meet the WHO target product profile for an optimal "one-and-done" LAI TPT clinical regimen.

## INTRODUCTION

Identification of latent *Mycobacterium tuberculosis* (TB) infection and provision of tuberculosis preventive therapy (TPT) is essential for global TB control.^1^ TPT is recommended for those at high risk of infection and progression to active TB disease.^2^ The World Health Organization (WHO) recommends six TPT regimens; five consist of isoniazid and/or a rifamycin (rifampin or rifapentine) and are given for durations of 1-9 months.^2^ Despite established efficacy and higher treatment completion rates for shorter rifamycin-based regimens, estimated global TPT coverage (21-56% in 2023) remains well below the United Nations High-Level Meeting on TB target of 90%.^3^ WHO recommends 6 months of levofloxacin for contacts of rifamycin-resistant (RR) or multidrug-resistant (MDR) TB, but this regimen is associated with low (68%) completion rates.^4,5^ Efficacious TPT regimens with high completion rates are necessary to treat a sufficient number of the estimated two billion people living with TB infection and meet elimination goals.^2,3^ Additionally, a clear unmet need exists for short-course rifamycin-free regimens for RR/MDR-TB contacts, or where rifamycin-based regimens are inappropriate due to drug-drug interactions or intolerance. The first-in-class diarylquinoline bedaquiline, approved by the United States Food and Drug Administration (FDA) and the European Medicines Agency for oral treatment of MDR-TB disease has demonstrated bactericidal activity in TPT mouse models,^6,7^ and is being evaluated in a 1-month daily oral regimen for TPT in the BREACH-TB trial.^8^

Long-acting drug delivery offers the potential to improve uptake and completion of TPT regimens,^9–11^ with surveys indicating provider and community group support for long-acting injectable (LAI) medicines.^12^ Several target product profiles have been proposed, with ideal long-acting TPT products providing “one-and-done” options via a single point-of-care administration.^9,10,13^ An LAI formulation of bedaquiline demonstrated bactericidal activity after intramuscular (IM) injection in TPT mouse models,^6,7^ and is currently in a phase 1 study for pharmacokinetics (PK), safety, and tolerability,^14,15^ providing a precedent for LAI-based TPT.

LAIs require high drug concentrations to achieve efficacious exposures within tolerable injection volumes.^9,16,17^ The bedaquiline LAI phase 1 study tested doses requiring up to two 5 mL IM injections, with preliminary reports indicating mostly grade 1-2 injection site reactions of pain or tenderness.^14,15,18^ The majority of participants reportedly expressed a preference for LAI over guideline-recommended oral 1-3 month isoniazid-rifapentine regimens.^18^ A phase 1 lenacapavir LAI study used similar IM injection volumes.^19^ Both studies reported pain as manageable with ice, analgesics, and/or counseling, but clearly lower volumes are desirable. Achieving this necessitates potent drugs, low systemic clearances, and slow depot site release rates, with ongoing developments of LAI formulations using increasingly potent diarylquinolines.^20,21^

Here, we present a preclinical characterization of novel LAI formulations of sorfequiline (TBAJ-876), a second-generation diarylquinoline. Sorfequiline is an order of magnitude more potent than bedaquiline *in vitro* and in mouse infection models,^22,23^ carries a lower risk of QT prolongation,^24,25^ and is currently being evaluated in a phase 2 trial in oral form to replace bedaquiline in the FDA-approved BPaL (bedaquiline-pretomanid-linezolid) regimen for treatment of TB disease.^26,27^

## METHODS

Detailed methods are provided in the Online Supplement.

### Sorfequiline LAI formulation

Sorfequiline tartrate was precipitated from a 1:1 mixture of acetone and methanol into an aqueous polyvinyl pyrrolidone (PVP) solution with ultrasound sonication. The suspended product was spray dried yielding a powdered sample.^28^ Three formulations containing 80 wt% sorfequiline utilized PVP K12PF (Formulation A), PVP C15 (Formulation B), or PVP K17PF (Formulation C).

### Mycobacterial strains

The live immunization strain rBCG30, provided by Marcus A. Horwitz,^29^ and *M. tuberculosis* H37Rv, American Type Culture Collection 27294, were each mouse passaged and frozen in aliquots.

### *In vivo* experiments

Adult (6-8 weeks) female BALB/c mice from Charles River (BALB/cAnNCrl, strain code 028) were used in all experiments. Mice were housed in individually ventilated cages (≤5 mice/cage) with sterile shredded paper for bedding (22-24°C; 12-hour light/12-hour dark cycle) and *ad libitum* access to food/water. The paucibacillary mouse model of TPT utilized immunization with rBCG30 prior to TB infection to establish a stable, low-burden TB lung infection suitable for testing TPT regimens.^6,7,30–32^ In experiments that included oral regimens, administration was by gavage 5 days/week. In experiments that included sorfequiline LAI regimens, IM injections (≤50 µL) were given in the right hind thigh. All PK sampling utilized in-life mandibular bleeding.^32^ For each experiment, a detailed study scheme including animal numbers is provided in the Online Supplement, and individual mouse data are provided in **Supplementary Data S1** (PK and TB data) and **Data S2** (rBCG30 data). Methods used for PK analyses are included in the Online Supplement.

## RESULTS

### Sorfequiline target plasma exposure estimation

No established PK/PD parameters exist to inform selection of sorfequiline target exposures for TPT. The minimum inhibitory concentrations (MICs) of sorfequiline and its major active M3 metabolite against *M. tuberculosis* H37Rv were determined to be 3.9 ng/mL and 16 ng/mL, respectively. While it is reasonable to assume that drug concentrations at the site of infection must be above the MIC to provide effective TPT, MICs are indices of *in vitro* growth inhibition determined against actively multiplying bacteria in nutrient-rich conditions, while TPT is indicated for smaller, slowly multiplying or non-multiplying bacterial populations under more stringent conditions. We previously demonstrated that approaches to target determination should utilize parameters derived or confirmed from *in vivo* efficacy studies.^7,16,32^ Therefore, based on modelling work used to estimate TPT target exposures for LAI formulations of other drugs including bedaquiline,^16^ as well as previous work in which data generated with orally-dosed drugs were used to estimate TPT target exposures for rifamycin LAI formulations,^32^ we designed a supporting experiment using orally-dosed sorfequiline to establish exposure-activity relationships in a paucibacillary mouse infection model suitable for evaluating TPT regimens.^30,31,33^

We tested three oral doses of sorfequiline, centered on a dose of 3 mg/kg/day, previously shown to be as bactericidal as bedaquiline 25 mg/kg/day (the established human-equivalent dose) in mouse models of TB disease,^22,34^ with bracketing doses of 1 and 10 mg/kg/day (experiment scheme in **Table S1**). After four weeks of treatment, bactericidal activity was observed across all oral doses, with the mean *M. tuberculosis* lung load decreasing by 0.97, 1.15, and 1.23 log_10_ colony-forming units (CFU) in mice that received sorfequiline at 1, 3, and 10 mg/kg/day, respectively, compared to the lung load in untreated mice (**Fig. S1A**). Sorfequiline and M3 plasma concentrations were measured during the third week of treatment. Plasma minimum concentrations (C_min_, 0 h) were proportional to dose, averaging 7.8, 37.8, and 126.7 ng/mL in mice receiving 1, 3, and 10 mg/kg sorfequiline orally, respectively. Plasma concentrations across all groups were approximately 2.5 times higher 3 h after dosing; for all dose levels, the plasma concentration of the M3 metabolite was approximately 4 times greater than the concentration of the parent moiety at both sampling time points (**Fig. S1B**). Measurement of M3 concentrations in mice is important for interpretation of efficacy data from mouse models and downstream translation to human studies, as M3 concentrations are much higher in mice relative to humans.^24,34^

The observed data aligned with previous data generated with orally-dosed sorfequiline in uninfected mice and in mouse models of active TB disease, but we were surprised by the limited dose-response effect observed and the inferior activity of sorfequiline at 10 mg/kg compared to the 1HP control regimen, when prior studies have shown bedaquiline at 25 mg/kg for 4 weeks or 32 mg/kg for 2 weeks has bactericidal activity similar to 1HP at 4 weeks.^6,30^ Therefore, a second experiment was performed, this time evaluating a wider dose-response and longer treatment duration, and including bedaquiline and another second-generation diarylquinoline, TBAJ-587, as comparators (scheme in **Table S2**). A distinct dose-response effect was observed, whereby sorfequiline at 10 mg/kg, as well as bedaquiline and TBAJ-587 at 25 mg/kg, had bactericidal activity similar to the 1HP control at 4 weeks (**Fig. S2A**), and 6 weeks of sorfequiline at 3 mg/kg demonstrated a significant bactericidal effect comparable to 1HP at 4 weeks (**Fig. S2B**).

Seeking plasma exposures above those observed with oral dosing of sorfequiline at 1 mg/kg (average C_min_ and C_3h_ of 7.8 and 18.7 ng/mL, respectively) and considering that the sorfequiline plasma concentration associated with 50% of its maximal effect (EC_50_) is 36.8 ng/mL when combined with pretomanid-linezolid (PaL) in our BALB/c mouse model of TB disease,^35^ we selected 35 ng/mL (with an associated M3 concentration approximately 4-fold higher) as a provisional target concentration above which bactericidal activity would be expected.

### Initial PK assessment of sorfequiline LAI formulations in mice

With a provisional exposure target established, PK assessment of three candidate sorfequiline formulations was conducted in uninfected mice after a single 50 µL injection of either the maximum syringeable concentration, or one-half or one-quarter of the maximum (∼1000, 500, and 250 mg/kg, respectively). Sorfequiline and M3 plasma concentrations were measured up to 12 weeks post-injection (scheme in **Table S3**). Mice that received the highest dose of formulation A required euthanasia according to protocol 3 days post-injection due to disuse of the injected leg. All other formulations/doses were well-tolerated.

PK exposure profiles consistent with long-acting release after administration were observed. Mean sorfequiline plasma concentrations were maintained above or approximating the provisional target C_min_ (35 ng/mL) ≥8 weeks post-injection across the study; at 12 weeks post- injection, mean plasma concentrations remained above or approximating the MIC of 3.9 ng/mL (**Fig. 1**; **Fig. S4**). To account for the contribution of the active M3 metabolite to the observed efficacy and enable the downstream translation of these non-clinical data to derive robust human plasma target exposures for sorfequiline LAIs, the “total combined equivalent” of sorfequiline concentration was calculated as a weighted sum of the sorfequiline and M3 plasma concentrations on the basis of M3 having a 4.19x higher molar MIC compared to sorfequiline (**Fig. 1**; calculation provided in Online Supplement).

**Figure 1.**
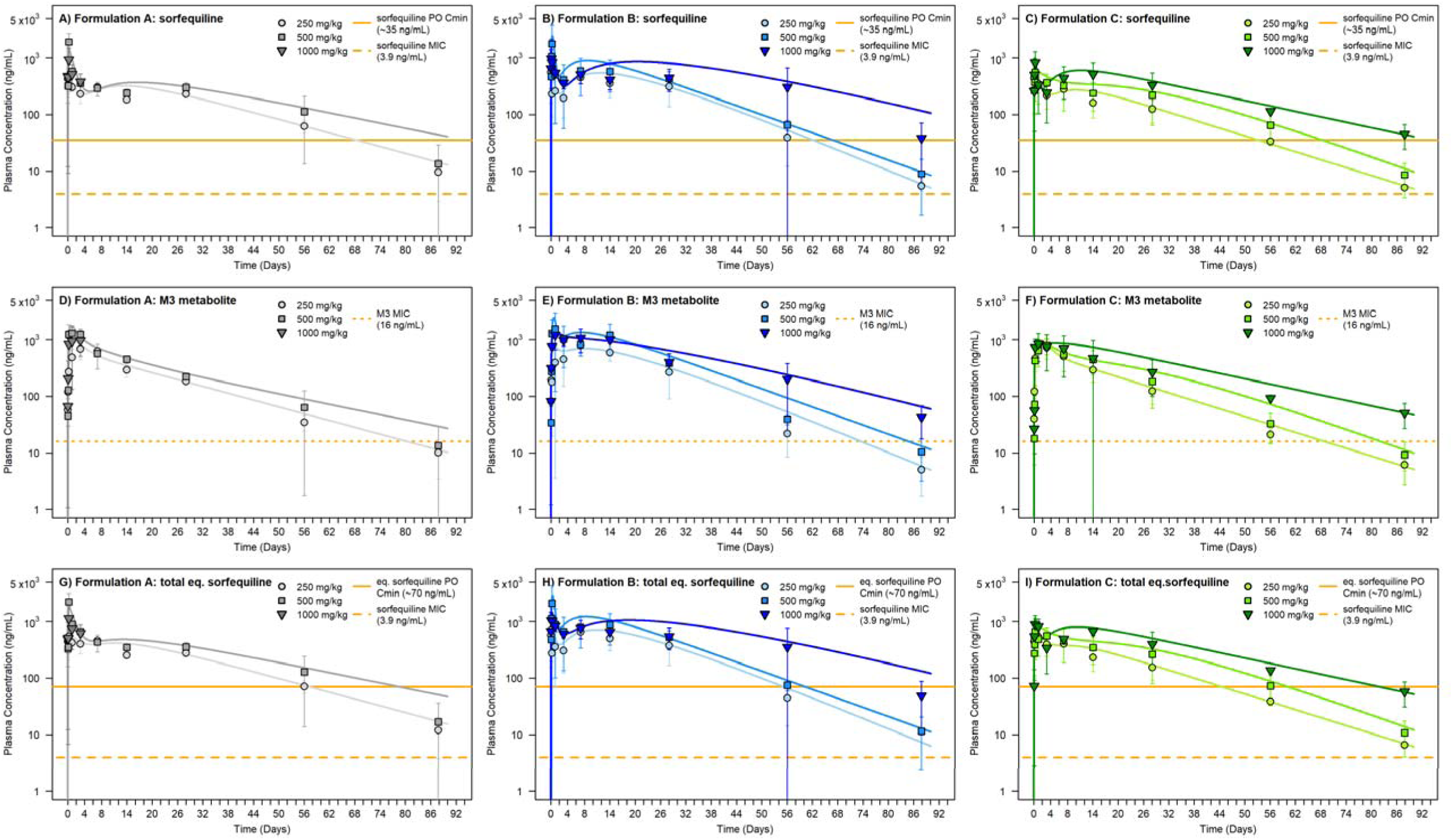
Plasma concentrations of sorfequiline (A-C), its major M3 metabolite (D-F), and the combined total equivalent (eq.) of sorfequiline (G-I) following a single IM injection of sorfequiline LAI formulations A, B, and C, each at three dose levels in uninfected female BALB/c mice, overlaid with simulation lines from compartmental PK model fittings to the profiles. The study scheme and sampling time points are presented in **Table S3** in the Online Supplement, and compartmental PK model schematic and parameter estimates in **Fig. S3** and **Table S4**, respectively. Data points represent the mean values and error bars represent standard deviation (n = 3 samples per time point). Plots are illustrated with reference lines of provisional targets or oral C_min_ and the MIC for sorfequiline parent, M3, or combined total equivalent of sorfequiline. Mice that received the highest dose for formulation A reached a humane endpoint 72 hours after injection and were removed from the study. Raw data are graphed in **Fig. S4**. All individual mouse data are provided in **Supplementary Data S1C**.

Non-compartmental summary PK parameters are presented in **Table 1**. Terminal phase sorfequiline half-lives of ∼10-20 days were observed for the LAI candidate formulations, far exceeding the terminal half-life following intravenous administration of 27 hours (TB Alliance NCT04493671 trial protocol;^36^ data on file). This indicates “flip-flop” PK,^37^ where observed half-lives reflect the slow release rates from a depot rather than disposition elimination based on drug clearance and volume of distribution. This was also reflected in compartmental PK analyses of sorfequiline and M3 plasma exposure, where observed profiles were adequately described with one-compartment disposition models, and “flip-flop” PK masking any more complex underlying disposition.^38^

**Table 1:** Table of PK parameters*^a^* for sorfequiline and its major M3 metabolite in uninfected and *M. tuberculosis* (Mtb)-infected mice.

| Sorfequiline LAI group<sup>b</sup> | Analyte | C<sub>max</sub><br>(ng/mL) | t<sub>max</sub><br>(h) | t<sub>1/2</sub><br>(days) | AUC<sub>0-last</sub><br>(µg·h/ml) | AUC<sub>0-inf</sub><br>(µg·h/ml) |
| --- | --- | --- | --- | --- | --- | --- |
| Formulation A 250 mg/kg<br>(uninfected mice) | Sorfequiline | 476 | 6 | 17 | 285 | 290 |
|  | M3 | 686 | 72 | 14 | 348 | 353 |
| Formulation A 500 mg/kg<br>(uninfected mice) | Sorfequiline | 1919 | 6 | 19 | 403 | 412 |
|  | M3 | 1298 | 24 | 15 | 498 | 505 |
| Formulation B 250 mg/kg<br>(uninfected mice) | Sorfequiline | 660 | 3 | 12 | 364 | 366 |
|  | M3 | 798 | 168 | 10 | 454 | 456 |
| Formulation B 500 mg/kg<br>(uninfected mice) | Sorfequiline | 1778 | 6 | 13 | 562 | 566 |
|  | M3 | 1550 | 24 | 11 | 778 | 782 |
| Formulation B 1000 mg/kg<br>(uninfected mice) | Sorfequiline | 975 | 3 | 24 | 686 | 717 |
|  | M3 | 1217 | 24 | 17 | 879 | 905 |
| Formulation C 250 mg/kg<br>(uninfected mice) | Sorfequiline | 514 | 3 | 14 | 200 | 203 |
|  | M3 | 753 | 24 | 13 | 303 | 305 |
| Formulation C 500 mg/kg<br>(uninfected mice) | Sorfequiline | 559 | 6 | 16 | 308 | 313 |
|  | M3 | 791 | 72 | 13 | 385 | 389 |
| Formulation C 1000 mg/kg<br>(uninfected mice) | Sorfequiline | 831 | 6 | 23 | 496 | 531 |
|  | M3 | 852 | 24 | 21 | 524 | 561 |
| Formulation A 125 mg/kg<br>(Mtb-infected mice) | Sorfequiline | 1017 | 2 | 11 | 221 | 222 |
|  | M3 | 285 | 168 | 11 | 139 | 140 |
| Formulation B 62.5 mg/kg<br>(Mtb-infected mice) | Sorfequiline | 420 | 24 | 7 | 144 | 147 |
|  | M3 | 283 | 72 | 8 | 103 | 105 |
| Formulation B 125 mg/kg<br>(Mtb-infected mice) | Sorfequiline | 475 | 168 | 11 | 218 | 219 |
|  | M3 | 306 | 168 | 12 | 147 | 148 |
| Formulation B 250 mg/kg<br>(Mtb-infected mice) | Sorfequiline | 924 | 2 | 11 | 413 | 416 |
|  | M3 | 517 | 336 | 12 | 281 | 285 |
| Formulation C 125 mg/kg<br>(Mtb-infected mice) | Sorfequiline | 710 | 2 | 10 | 202 | 203 |
|  | M3 | 269 | 168 | 11 | 123 | 124 |

<sup>a</sup>PK parameters: C<sub>max</sub>, t<sub>max</sub>, maximum plasma concentration and time of occurrence; t<sub>1/2</sub>, half-life of terminal exponential phase of profile; AUC<sub>0-last</sub>, area under plasma concentration-time profile curve from 0 to last observed timepoint; AUC<sub>0-inf</sub>, area under plasma concentration-time profile curve from 0 extrapolated to infinity.
<sup>b</sup>Uninfected mice from initial PK study with sorfequiline LAI formulations; Mtb-infected mice from PK/PD study with sorfequiline LAI formulations in a mouse model of TPT.

Simultaneous compartmental PK analysis of sorfequiline and M3 found sorfequiline profiles were adequately described with two first-order release input processes, one fraction releasing faster and driving the initial maximum plasma concentration (C_max_), and one releasing slower and contributing to the majority of the area under the plasma concentration-time profile curve (AUC) (**Fig. 1**). M3 PK profiles were only adequately described when M3 production was modelled to occur both as a result of plasma clearance of sorfequiline and as a fraction produced and absorbed directly at the administration site via a theorized local first pass effect (**Fig. S3**; **Table S4** in the Online Supplement). Production of M3 as a result of a first pass effect has previously been modelled to occur with oral dosing of sorfequiline.^38^ Unlike for the oral route, the precise mechanism(s) for first pass metabolism and extraction following IM injection remain uncertain but may be due to low-level enzymatic drug metabolism in muscle tissue, absorption into the lymph, or immune cell clearance. Calculations indicated an observed bioavailability of <100% at the higher 500 and 1000 mg/kg doses (**Table S4**), consistent with depot site metabolism.

### Sorfequiline LAI PK/PD evaluation

The promising results from the PK study supported advancing to a proof-of-concept efficacy study using the validated paucibacillary mouse model of TPT. As data from the initial PK study indicated that formulation B exhibited slightly higher plasma exposures compared to formulations A and C at similar dosing levels (**Fig. 1**; **Table 1**), formulation B was selected for dose-ranging PK/PD assessment in the model. A dose of 250 mg/kg was selected for efficacy evaluations based on its ability to maintain concentrations at or above the provisional exposure target for 8 weeks in mice as well as newly available phase 1 human PK data suggesting that the average daily plasma AUC and C_max_ based on the sum of active moieties (*i.e.*, total equivalent sorfequiline) were roughly similar to and lower than, respectively, values observed in humans receiving oral sorfequiline at 200 mg/day for 14 days, which was found to be safe and well- tolerated.^24^ We also tested the maximum dose of 250 mg/kg administered as a single injection or administered as two injections, each 125 mg/kg, spaced 2 weeks apart.

Other doses were selected as fractions of the maximum dose in two steps to evaluate efficacy of single injection doses of 125 mg/kg (½ max) and 62.5 mg/kg (¼ max). A head-to-head comparison of the three different sorfequiline LAI formulations was incorporated by administering formulations A and C at 125 mg/kg in a single injection. We evaluated the efficacy of each of these LAI formulations in the mouse model of TPT up to 12 weeks from start of treatment. Bactericidal activity was compared to oral control regimens (1HP and bedaquiline 25 mg/kg/day [total bedaquiline dose 500 mg/kg]) as well as two sorfequiline oral comparator regimens: 6.25 mg/kg/day (total dose 125 mg/kg) and 12.5 mg/kg/day (total dose 250 mg/kg).

All oral regimens were administered 5 days per week for 4 weeks. The complete study scheme is presented in **Table S5** (treatment groups and time points for bactericidal activity assessment) and **Table S6** (PK sampling scheme).

All LAI and oral sorfequiline regimens exhibited bactericidal activity that was superior to 1HP (p<0.0001), based on between-group comparisons of mean CFU counts (**Fig. 2A**; **Table S7**); and sorfequiline regimens had similar activity when the same total sorfequiline dose was given (e.g., 125 mg/kg IM vs. 6.25 mg/kg oral). Sorfequiline LAI formulations A, B, and C had similar bactericidal activity when dosed at 125 mg/kg via a single IM injection; and activity was dose-dependent for formulation B (**Fig. 2B**; **Table S7**). Administration of formulation B in a single 250 mg/kg injection or two injections of 125 mg/kg, spaced 2 weeks apart, resulted in similar bactericidal activity (**Table S7**). Among all mice that received a sorfequiline LAI formulation, only six had culture-positive lungs at Week 12, and none of these cultured isolates exhibited evidence of decreased sensitivity to diarylquinolines (**Table S8**). All sorfequiline LAI regimens exhibited a time course consistent with bactericidal activity while plasma sorfequiline exposures exceeded the provisional target C_min_ of 35 ng/mL (**Fig. 2C-D** [sorfequiline parent]; **Fig. 2E-F** [M3 metabolite]; **Fig. 2G-H** [total sorfequiline equivalent]); these data support the selected provisional target. In all, 23 of 25 (92%) mice receiving at least 125 mg/kg of any sorfequiline LAI formulation were rendered culture-negative at 12 weeks. Plasma exposure (AUC) and half-life for sorfequiline and the M3 metabolite were generally dose-proportional and consistent with those seen in uninfected animals, although the ratio of sorfequiline AUC to M3 AUC was higher in the experiment using infected mice (**Table 1**, **Fig. 2**).

**Figure 2.**
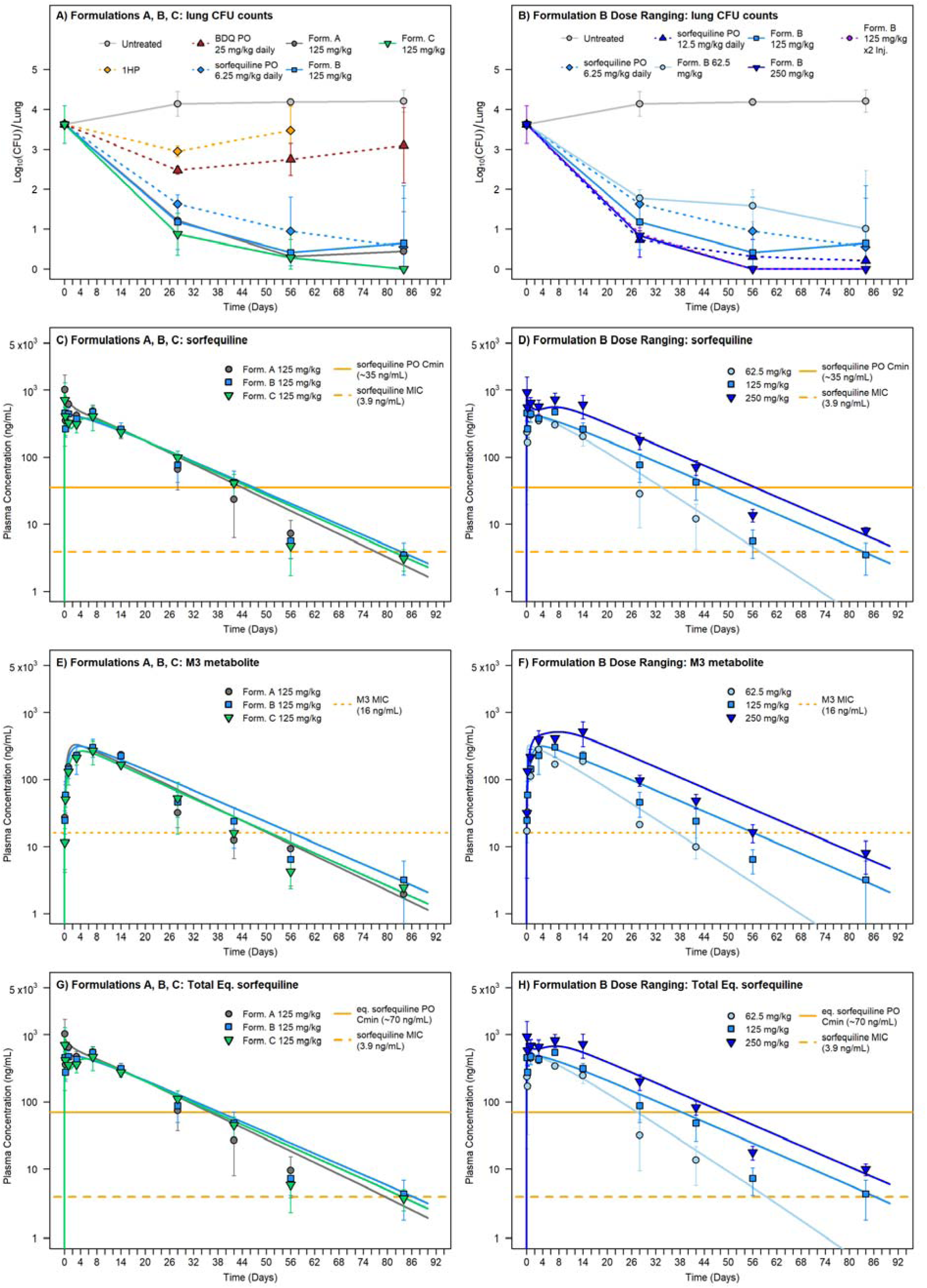
Lung CFU counts associated with treatment in the paucibacillary mouse model of TPT with a single injection of sorfequiline formulations A, B and C at 125 mg/kg (Panel A) and dose ranging of a single injection of sorfequiline formulation B at 62.5, 125, and 250 mg/kg, and 2 injections at 125 mg/kg separated by 2 weeks (Panel B), with reference control oral treatment profiles. CFU data points represent mean values, and error bars represent standard deviation (n = 3-5 mice per time point). Matching associated plasma concentration profiles in the TPT mouse model animals following these treatments, overlaid with simulated lines from the compartmental PK model fittings to the data, are presented for sorfequiline (panels C and D) its major M3 metabolite (panels E and F) and the combined total equivalent (Eq.) of sorfequiline (panels G and H). PK data points represent mean values, and error bars represent standard deviation (n = 3 samples per time point). PK plots are illustrated with reference lines of provisional targets of oral C_min_ and MIC of sorfequiline parent, M3, or combined total equivalent of sorfequiline. Control orally dosed (PO) treatments were given once daily, 5 days per week (Monday-Friday) for 4 weeks with doses as follows:1HP: isoniazid and rifapentine, each at 10 mg/kg. BDQ: bedaquiline 25 mg/kg. sorfequiline at 6.25 or 12.5 mg/kg. The study design and PK sampling schemes are provided in **Tables S5** and **S6**, respectively, in the Online Supplement. Raw data are graphed in **Fig. S5**. Summary CFU data for all study groups are presented in **Table S7**, and individual mouse data are provided in **Supplementary Data S1D**.

## DISCUSSION

We present proof-of-concept data for a highly efficacious TPT regimen from a single IM sorfequiline injection. All candidate formulations (A and C at 125 mg/kg; B at 62.5-250 mg/kg) showed significantly greater bactericidal activity than the shortest WHO-recommended oral TPT regimen (1HP), known to be non-inferior to 9-months of isoniazid among persons living with HIV in the BRIEF-TB trial.^33^ Importantly, 1HP has comparable efficacy to three other WHO-recommended regimens including 3HP (once-weekly high-dose isoniazid-rifapentine; 3 months), and superiority against daily isoniazid (6 months) in the paucibacillary mouse model.^30^ Therefore, a sorfequiline LAI may offer a one-shot option for TPT, if comparable exposures can be achieved clinically.

A short-course regimen suitable for TPT in people exposed to RR-TB is a major unmet need, as is a short-course TPT regimen for people unable to take a rifamycin due to drug-drug interactions or intolerance. Indeed, a sufficiently safe and effective rifamycin-sparing short-course regimen could be used as a “pan-TPT” regimen to treat individuals without knowledge of the isoniazid, rifamycin or fluoroquinolone susceptibility of the presumed infecting strain.

Diarylquinolines are the best candidates for such a regimen given their strong sterilizing activity, lower drug-drug interaction potential, and the key role of bedaquiline in RR-TB treatment.

Implementation experience of bedaquiline for TPT is very limited, yet promising. In one unpublished report, a 3-month regimen of oral bedaquiline was well-tolerated and safe, and associated with no incident TB disease after 2 years of follow-up among 41 participants exposed to index patients with (pre-)extensively drug-resistant TB.^39^ The randomized controlled BREACH-TB trial is evaluating efficacy, safety, and tolerability of oral bedaquiline for 1 month as TPT in close contacts of drug-susceptible or RR-TB or people living with HIV infection in areas with high TB transmission.^8^

Our results show similar efficacy of oral bedaquiline (1 month) and 1HP in the paucibacillary mouse model of TPT and further support the potential for diarylquinoline-based short-course regimens for TPT. Sorfequiline has superior *in vitro* and *in vivo* potency over bedaquiline,^22,23,34^ and results from the phase 2 NC-009 trial indicate replacement of daily bedaquiline (200 mg; BPaL regimen) with sorfequiline (100 mg daily) offers faster sputum culture conversion in participants with drug-susceptible TB.^27^ Our present results with oral sorfequiline doses producing plasma exposures comparable to the 100 mg daily dose in humans showing superior efficacy over bedaquiline and 1HP suggest that oral sorfequiline for durations shorter than one month may be effective as TPT.

Bedaquiline LAI formulations are in clinical development, and preliminary results of a phase 1 single ascending dose study in healthy volunteers were recently reported.^14,18^ A single 5 mL injection achieved bedaquiline target exposures predicted to be as efficacious as 3HP based on a translational model, with few adverse events beyond transient Grade 1 or 2 injection site pain/tenderness. Most participants stated a preference for the LAI over 1HP or 3HP regimens to complete TPT.

We have not compared sorfequiline LAI candidates against the bedaquiline LAI. However, we previously found that a single 160 mg/kg IM injection of a bedaquiline microsuspension LAI was less bactericidal than 1HP and approximately equivalent to 10 doses of oral bedaquiline at 32 mg/kg (320 mg/kg total dose) administered over a 12-day period.^6^ In contrast, a single sorfequiline 62.5 mg/kg IM injection (Formulation B) was significantly more bactericidal than 1HP and 20 doses of oral bedaquiline (25 mg/kg; 500 mg/kg total dose) administered over a 26- day period. In comparison to the control regimens (1HP and oral bedaquiline), sorfequiline LAIs are more active at lower doses than the bedaquiline LAI in this paucibacillary model. If sufficient exposure is replicated clinically, the potency advantage of sorfequiline coupled with high drug loading and syringeability of our formulations should translate to superior efficacy and/or tolerability (smaller injection volumes) compared to the bedaquiline LAI formulation currently in phase 1. Importantly, the LAI formulations presented here have high potential for commercial translation with simple manufacturability and no highly specialized equipment. The dry powdered product aims to avoid cold chains often employed in liquid suspension storage. The excipients used are well studied and tolerated and are readily available in pharmaceutical grade versions.

Although murine paucibacillary models have informed downstream clinical development of efficacious short-course TPT regimens and ongoing evaluations of newer candidate regimens, estimation of drug exposures required for clinical TPT efficacy from preclinical data remains challenging. Little is known about the exposures required for minimum/optimum clinical efficacy of any TPT regimen, and current regimens use the same doses of isoniazid, rifampicin, rifapentine, and levofloxacin proven effective in treatment of active TB disease, although it is notable that the 3HP regimen more closely resembles the labelled dosing and schedule for the continuation phase of TB treatment.^40,41^ With the possible exception of isoniazid monotherapy, durations required for any TPT regimen have not been meaningfully determined. Therefore, exposures required for efficacious LAI TPT candidate regimens is unclear, especially when coupled to drugs with no established oral TPT regimen. Differences in the shapes of PK profiles produced by oral administration compared to their LAI formulations make it difficult to translate effective oral exposure into predictions for effective LAI exposure. The paucibacillary mouse model can provide useful information to bridge this knowledge gap by establishing *in vivo* exposure-activity relationships in a model validated for translational and predictive value. To appropriately interpret and translate the sorfequiline (oral and LAI) efficacy data generated in mice, the more rapid metabolism, relative to humans, of sorfequiline to the less potent M3 metabolite must be accounted for (M3:parent ratio is much higher and thus the M3 contributes to the observed efficacy in mice). Therefore, based on work dealing with a similar issue with bedaquiline and its M2 metabolite,^42–44^ we measured M3 concentrations and calculated total equivalent sorfequiline exposure to include in our PK/PD analyses and to inform future mouse-to-human translation of sorfequiline PK/PD data. However, given the robust exposure and efficacy data observed with our sorfequiline LAI formulations, these (current) translational challenges do not change our overall conclusions. The demonstrated superior efficacy of sorfequiline LAI formulations to 1HP indicates their potential to meet the WHO target product profile for an optimal LAI TPT regimen^45^ and, as such, warrant their further development.

## Author contributions

N.C.A., H.P., A.O., J.S., E.L.N.: conceptualized the study. E.L.N., A.O., N.F.: led funding acquisition for the study. J.J.H., N.C.A., S.R., E.L.N.: drafted the manuscript. H.P., N.C.A.: led data analysis and visualization. N.C.A, H.P., E.L.N., A.O.: contributed to data curation and analysis. J.J.H, S.R.: preparation of long-acting sorfequiline formulations. S.Y.L., N.C.A., E.L.N.: involved in data collection and study investigation. All authors critically reviewed drafts, supported data interpretation, and approved the manuscript for submission.

## Data availability statement

Experimental data are provided in the online data supplement. R code scripts are available upon request.

## Ethical approval statement

Experiments using mice were conducted at the Johns Hopkins University Center for TB Research under protocols approved by the Institutional Animal Care and Use Committee of the Johns Hopkins University.

## Artificial intelligence (AI)-assisted technology

AI-assisted technology was not used in the production of the submitted work.

