## Supplemental Methods, Tables, and Figures for "One-Dose Efficacy of Long-Acting Sorfequiline in a Mouse Model of Tuberculosis Preventive Therapy"

**ONLINE SUPPLEMENT**

**Abbreviations Used**	**2**

**Supplementary Methods**	**3**

Sorfequiline LAI formulations	3

Media preparation for mycobacterial growth, MIC assays, and CFU determination	3

MIC assays	3

Preparation of drugs/compounds for oral administration to mice	3

Preparation of sorfequiline LAI formulations for administration to mice	4

Mouse lung processing for CFU determination	4

Assessment of DARQ susceptibility in mouse lung homogenates	4

PK sampling and processing	4

Bioanalysis to determine sorfequiline and M3 metabolite plasma concentrations	4

PK analyses	5

**Supplementary Tables**	**6**

**Table S1.** Scheme of PK/PD study with oral dose-ranging of sorfequiline in a mouse model of TPT	6

**Table S2.** Scheme of PD study with oral dosing of DARQs in a mouse model of TPT	7

**Table S3.** Scheme of PK study with sorfequiline LAI formulations in uninfected mice	8

**Table S4.** Table of parameter estimates for sorfequiline PK model	9

**Table S5.** Scheme of PK/PD study with sorfequiline LAI formulations in a mouse model of TPT	10

**Table S6.** PK sampling scheme for PK/PD study with sorfequiline LAI formulations	11

**Table S7.** Summary bactericidal activity (lung CFU) data for PK/PD study with sorfequiline LAI formulations	12

**Table S8.** Assessment of DARQ susceptibility in Week 12 lung homogenates from PK/PD study	13

**Supplementary Figures**	**14**

**Figure S1.** PK/PD data from study with oral dose-ranging of sorfequiline in a mouse model of TPT	14

**Figure S2.** *M. tuberculosis* H37Rv lung CFU data from study with oral dose-ranging of DARQs in a mouse model of TPT	15

**Figure S3.** Schematic and differential equations for the sorfequiline compartmental PK model	16

**Figure S4.** PK data from PK study with sorfequiline LAI formulations in uninfected mice	17

**Figure S5.** Lung CFU and PK data from PK/PD study with sorfequiline formulations in a mouse model of TPT	18

**References**	**19**

**Supplementary Individual Mouse PK/PD Data**	**File S1**

Individual mouse data from PK/PD study with oral dosing of sorfequiline	Tab S1A

Individual mouse data from PD study with oral dosing of DARQs	Tab S1B

Individual mouse data from PK study with sorfequiline LAI injectable formulations	Tab S1C

Individual mouse data from PK/PD study with sorfequiline LAI injectable formulations	Tab S1D

**Supplementary Individual Mouse rBCG30 CFU Data**	**File S2**

Individual mouse data from PK/PD study with oral dosing of sorfequiline	Tab S2A

Individual mouse data from PD study with oral dosing of DARQs	Tab S2B

Individual mouse data from PK/PD study with sorfequiline LAI injectable formulations	Tab S2C

**ABBREVIATIONS USED**

| API | active pharmaceutical ingredient |
| --- | --- |
| BDQ | bedaquiline |
| CFU | colony-forming unit(s) |
| CRO | contract research organization |
| DARQ | diarylquinoline |
| DMSO | dimethyl sulfoxide |
| h | hour(s) |
| H | isoniazid |
| IM | intramuscular |
| LAI | long-acting injectable |
| MIC | minimum inhibitory concentration |
| Mtb | *Mycobacterium tuberculosis* |
| OADC | oleic acid-albumin-dextrose-catalase |
| P | rifapentine |
| PD | pharmacodynamic(s) |
| PK | pharmacokinetic(s) |
| PO | *per os* (by mouth) |
| TB | tuberculosis |
| TPT | tuberculosis preventive therapy |
| w | weeks(s) |

**SUPPLEMENTARY METHODS**

**Sorfequiline LAI formulations**

The sorfequiline LAI formulations have been previously partially reported at the following scientific meetings: Conference on Retroviruses and Opportunistic Infections (CROI) 2024 in Denver, Colorado; Centre of Excellence for Long-acting Therapeutics (CELT) Global Health inaugural meeting 2024 in Liverpool, UK; American Association of Pharmaceutical Scientist (AAPS) 2025 PHARMSCI 360 meeting in San Antonio, Texas; and SCI Formulation Forum 2026 in London, UK. Further details regarding the development and characterization of the formulations will be presented in a separate publication.

**Media preparation for mycobacterial growth MIC assays, and CFU determination**

The standard liquid growth medium for both mycobacterial strains (rBCG30 and *M. tuberculosis* H37Rv) was Middlebrook 7H9 broth supplemented with 10% (vol/vol) Middlebrook OADC enrichment, 0.5% (vol/vol) glycerol, and 0.1% (vol/vol) Tween 80. All solid medium was based on *Mycobacteria* 7H11 agar supplemented with 10% (vol/vol) OADC and 0.5% (vol/vol) glycerol. For MIC assays, sorfequiline and M3 metabolite were dissolved in DMSO and added to the agar for a final DMSO concentration of 0.125% (vol/vol), with DMSO alone added to agar for growth controls samples. For CFU determination, mouse lung homogenates and their cognate 10-fold dilutions were cultured on 7H11 agar rendered selective for mycobacteria by the addition of 50 mg/L carbenicillin, 10 mg/L polymyxin B, 20 mg/L trimethoprim, and 50 mg/L cycloheximide. Selective 7H11 agar was also supplemented with 0.4% (wt/vol) activated charcoal to detect and limit the effects of drug carryover in lung homogenates. To differentiate rBCG30 and *M. tuberculosis* CFUs, selective agar was further supplemented with either 40 mg/L hygromycin B (selective for *M. bovis* rBCG30 but not *M. tuberculosis*) or 200 mg/L 2-thiophenecarboxylic acid hydrazide (selective for *M. tuberculosis* but not *M. bovis*). Difco Middlebrook 7H9 broth powder, Difco *Mycobacteria* 7H11 agar powder, and BBL Middlebrook OADC enrichment were manufactured by Becton, Dickinson and Company. Glycerol and Tween 80 were purchased from ThermoFisher Scientific, and activated charcoal was manufactured by J.T. Baker. All selective drugs and DMSO were purchased from Sigma-Aldrich/Millipore-Sigma. Sorfequiline and M3 powders used in MIC assays were provided by the TB Alliance.

**MIC assays**

Sorfequiline and M3 metabolite MICs were determined for *M. tuberculosis* H37Rv using the agar proportion method.^1^ Sorfequiline or M3 powders dissolved in DMSO were diluted in agar at test concentrations ranging from 0.4-250 ng/mL (sorfequiline) and 8-500 ng/mL (M3). Data represent the geometric mean MIC determined from 3 replicates (sorfequiline) or 2 replicates (M3).

**Preparation of oral drugs/compounds for administration to mice**

Dosing (mg/kg) was based on an average mouse body mass of 20 g, with an oral administration volume of 0.2 mL. Isoniazid powder was purchased from Sigma-Aldrich/Millipore-Sigma and was dissolved in distilled water for administration to mice. Rifapentine tablets (PRIFTIN) were purchased from the Johns Hopkins Hospital Pharmacy and were crushed and suspended in 0.05% (wt API/vol) agarose solution. Bedaquiline fumarate powder was purchased from Biosynth; sorfequiline tartrate and TBAJ-587 powders were provided by the TB Alliance, and each of these agents was dissolved in 20% (wt/vol) hydroxypropyl-cyclodextrin solution acidified with 1.5% (vol/vol) 1N HCl. Invitrogen UltraPure Agarose powder was purchased form ThermoFisher Scientific, and hydroxypropyl-cyclodextrin and HCl were purchased from Sigma-Aldrich/Millipore-Sigma.

**Preparation of sorfequiline LAI formulations for administration to mice**

Sorfequiline LAI formulations A, B, and C, each at 80% API loading, were administered to mice in two studies. Prior to each study, LAI powders were aliquoted into glass vials (one vial per formulation/dose) at the University of Liverpool and then shipped at ambient temperature to the Johns Hopkins Center for TB Research, where they were stored inside of a desiccator at ambient temperature until use. About 1 hour prior to injection, the LAI powders were suspended in sterile water and dispersed by vortexing 30-60 seconds. The suspensions were vortexed again immediately prior to syringe loading and injection using a 25G needle.

**Mouse lung processing for CFU determination**

After killing via cervical dislocation under inhalation anesthesia with isoflurane, the entire lung was dissected from each mouse and placed into a Precellys 7 mL tissue grinding tube (CKMix50) containing 2.5 mL PBS. The lungs were then homogenized directly in these tubes using the Precellys Evolution touch, 7,200 rpm for 16 seconds. Ten-fold dilutions of lung homogenates were prepared with PBS (100 µL + 900 µL PBS), and 500 µL of lung homogenates and/or their cognate dilutions were cultured on appropriate agar plates. Detailed methods regarding reading the agar plates and calculation of total lung CFUs have been previously described.^2^

**Assessment of DARQ susceptibility in mouse lung homogenates**

In the sorfequiline LAI PK/PD experiment, mouse lung homogenates were also assessed for the presence of mycobacteria with decreased susceptibility to DARQs. Per established practice, BDQ was used as a representative DARQ to benchmark susceptibility in mycobacteria cultured from mouse lungs.^3^ At Week 12, one-fifth of the undiluted lung homogenate (0.5 mL) was directly cultured on plain 7H11 agar containing 0.125 mg/L BDQ. This agar did not contain activated charcoal in order to prevent adsorption of BDQ; this agar also did not contain HYG or TCH, and therefore mycobacterial CFUs could represent both Mtb and rBCG30. The number of CFU/lung cultured on BDQ-containing agar was then divided by the total number of CFU/lung as determined from lung homogenate plated on plain agar (i.e., agar without HYG or TCH) containing activated charcoal, and this ratio was used as a proxy for frequency of resistance to BDQ 0.125 mg/L. As this assessment was performed only at Week 12, the ratio determined in lung homogenates from DARQ-treated mice was compared to the ratio determined in the untreated mice at the same timepoint.

**PK sampling and processing**

At each sampling timepoint, 25-30 µL blood was collected by in-life mandibular bleeding into Becton-Dickinson vacutainer separation tubes with lithium heparin, and plasma was separated by centrifugation at 15,000 × *g* for 5 minutes at room temperature. Plasma was transferred using low-retention pipette tips into 1.5 mL O-ring screw-cap tubes and stored at -80°C.

**Bioanalysis to determine sorfequiline and M3 metabolite plasma concentrations**

Bioanalysis of the mouse plasma samples was conducted by the CRO Resolian (formerly Alliance Pharma and Drug Development Solutions). Per the requirements of the CRO, plasma samples were thawed prior to shipping, decontaminated with ACN (15 µL plasma + 30 µL ACN), refrozen, and then shipped on dry ice to the CRO for bioanalysis. Processing of the decontaminated mouse plasma and measurement of sorfequiline and its M3 metabolite concentrations were performed via CRO-developed and validated methods using high-performance liquid chromatography tandem mass spectrometry. The quantitative range for both sorfequiline and its M3 metabolite was 1.00-1000 ng/mL. Because the samples were diluted as part of the decontamination procedure, all measured concentrations were below the 1000 ng/mL standard.

**PK analyses**

Non-compartmental PK analysis and PK and simulations were performed in the R programming environment (v4.3).^4^ Compartmental PK models were fitted simultaneously to observed parent and metabolite data making use of the Pracma library,^5^ with parameter estimation by nonlinear regression using the “Isqnonlin” function for nonlinear least-squares optimization, with an objective function weighted by 1/(predicted value)^2. Data from all individual mice in any given data set were treated as a naïve pool^6^ rather than using an average value at a given time point. R code scripts used for analysis are available upon request.

“Total combined equivalent” sorfequiline concentration weighted sum was calculated as follows:

sorfequiline MIC = 3.9 ng/mL = 5.93 nM (molecular weight sorfequiline = 657)

M3 MIC = 16 ng/mL = 24.9 nM (molecular weight M3 = 643)

M3 MIC is 4.19x higher than sorfequiline parent, therefore (by crude approximation) M3 concentrations are 4.19x less efficacious than sorfequiline parent. A weighted sum of sorfequiline equivalent is therefore given by:

((molar concentration of M3)/4.19) + molar concentration of sorfequiline

Which can then be converted to ng/mL for sorfequiline by multiplying by the molecular weight (657).

This weighted sum calculation is also equivalent to:

(((concentration of M3)/(M3 MIC)) + ((concentration of sorfequiline)/(sorfequiline MIC))) x (sorfequiline MIC)

i.e., the sum of the individual moiety MIC ratios, multiplied by the MIC of sorfequiline.

**Table S1.** Scheme of PK/PD study with oral dose-ranging of sorfequiline in a paucibacillary mouse model of TPT. The original plan was to start treatment six weeks after the *M. tuberculosis* challenge infection; however, unforeseen logistical issues resulted in an extended period of 16 weeks between challenge infection and the start of treatment. We had previously demonstrated that extending the time between challenge infection and the start of treatment up to 22 weeks did not negatively impact performance of the model.^7^ Study timepoints are relative to the start of treatment on Day 0. 1HP (H_10-PO_P_10-PO_) indicates one month (4 weeks) of daily oral isoniazid (H) and rifapentine (P), both at 10 mg/kg. For the mice receiving sorfequiline oral dosing, PK sampling was conducted on Day 23 of this study, which corresponded to the 18^th^ dose of treatment (n = 3 mice per dosing group were sampled at two timepoints: just prior to (0 h) and 3 hours after dosing).

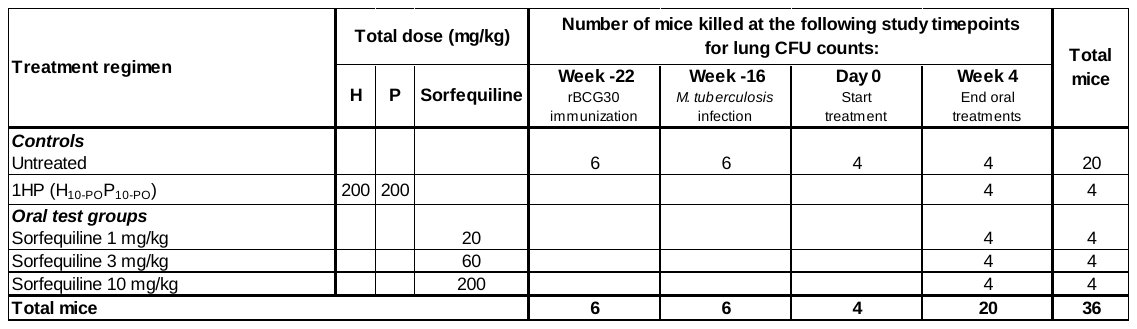

**Table S2.** Scheme of PD study with oral dose-ranging of DARQs bedaquiline, TBAJ-587, and sorfequiline in the paucibacillary mouse model of TPT. Study timepoints are relative to the start of treatment on Day 0. 1HP (H_10-PO_P_10-PO_) indicates one month (4 weeks) of daily oral isoniazid (H) and rifapentine (P), both at 10 mg/kg.

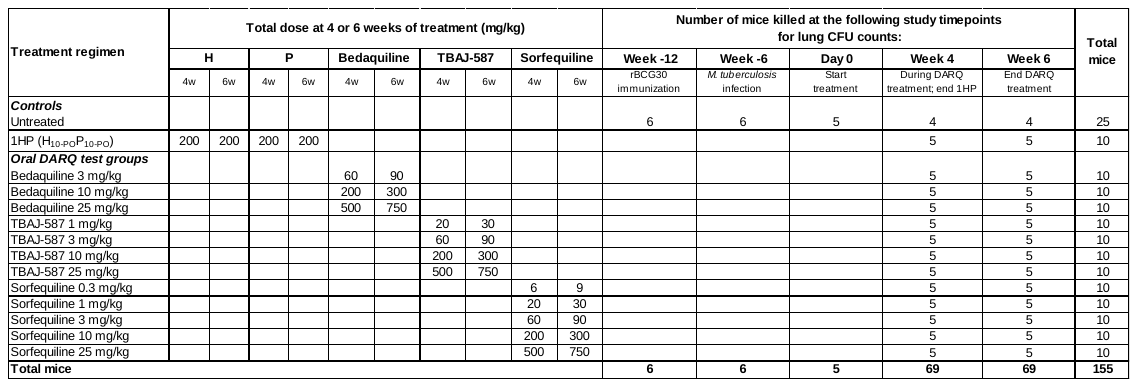

**Table S3.** Scheme of the initial PK study for sorfequiline LAI formulations A, B, and C in uninfected mice. At each timepoint post-intramuscular injection, blood was obtained by in-life mandibular bleeding from three mice (e.g., M1-M3; M4-M6, etc.), and shading in the table is used to identify the three-mouse cohorts for each formulation/dose sampled at each timepoint.

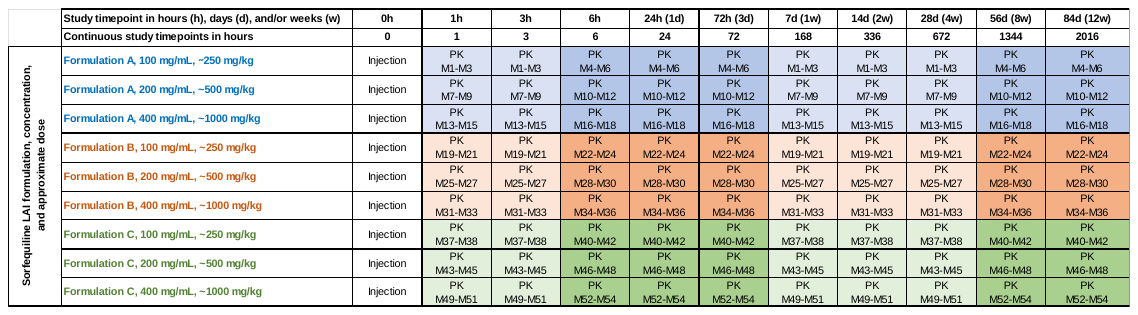

**Table S4.** Table of parameter estimates for PK model applied in simultaneous nonlinear regression fittings to plasma PK profiles for sorfequiline and its M3 metabolite. CL_par_: Clearance of parent sorfequiline; V_par_: volume of distribution of parent sorfequiline; KA1: first order release rate constant for fast release fraction; F2: fraction of dose assigned to slow release; KA2: first order release rate constant for slow release fraction; KT: first order rate constant for transit compartment delay of slow release fraction; CL_met_: Clearance of M3 metabolite V_met_: volume of distribution of M3 metabolite; FAM: fraction of F2 fraction assigned for release as M3 metabolite; F.app: apparent bioavailability (F % bioavailability calculated versus AUC_0-inf_ of 3.185 μg·h/mL for sorfequiline following an intravenous dose in mice of 2.5 mg/kg (TB Alliance NCT04493671 trial protocol;^8^ data on file). NCA: non-compartmental analysis; RSE: % relative standard error of estimate. See experimental scheme in **Table S3**. All individual mouse data are provided in **Supplementary Data S1C**.

| **Sorfequiline LAI group** | **CL_par_** | **V_par_** | **KA1** | **F2** | **KA2** | **KT** | **CL_met_** | **V_met_** | **FAM** |  | **F app.1** | **F app.2** |
| --- | --- | --- | --- | --- | --- | --- | --- | --- | --- | --- | --- | --- |
|  | **(L/h)** | **(L)** | **(h^-1^)** | **(%)** | **(h^-1^)** | **(h^-1^)** | **(L/h)** | **(L)** | **(%)** |  | **from NCA** | **from FAM est.** |
|  | **[RSE]** | **[RSE]** | **[RSE]** | **[RSE]** | **[RSE]** | **[RSE]** | **[RSE]** | **[RSE]** | **[RSE]** |  | **(%)** | **(%)** |
| Formulation A 250 mg/kg  uninfected mice | 0.010 | 0.57 | 735.3 | 94% | 0.0069 | 0.0019 | 0.01 | 0.2612 | 39% |  | 91% | 64% |
|  | [18.2] | [46.4] | [0.0] | [2.5] | [31.9] | [8.0] | [9.0] | [24.5] | [25.3] |  |  |  |
| Formulation A 500 mg/kg  uninfected mice | 0.012 | 0.42 | 2.10 | 92% | 0.0049 | 0.0015 | 0.02 | 0.1573 | 46% |  | 65% | 58% |
|  | [30.1] | [40.9] | [0.2] | [3.0] | [40.3] | [16.8] | [12.9] | [33.8] | [32.3] |  |  |  |
| Formulation B 250 mg/kg  uninfected mice | 0.008 | 0.17 | 3.52 | 97% | 0.0030 | 0.0050 | 0.01 | 0.0657 | 36% |  | 115% | 65% |
|  | [30.8] | [61.1] | [0.0] | [1.5] | [17.0] | [40.1] | [13.8] | [31.4] | [53.4] |  |  |  |
| Formulation B 500 mg/kg  uninfected mice | 0.011 | 0.08 | 0.13 | 94% | 0.0026 | 0.0087 | 0.01 | 0.0830 | 31% |  | 89% | 71% |
|  | [35.8] | [158.4] | [142.5] | [2.4] | [10.6] | [52.2] | [17.2] | [38.7] | [83.8] |  |  |  |
| Formulation B 1000 mg/kg  uninfected mice | 0.008 | 0.23 | 3.35 | 98% | 0.0021 | 0.0023 | 0.02 | 0.1389 | 55% |  | 56% | 46% |
|  | [170.3] | [175.8] | [0.6] | [3.0] | [129.4] | [139.0] | [12.6] | [56.7] | [141.2] |  |  |  |
| Formulation C 250 mg/kg  uninfected mice | 0.016 | 0.66 | 1.11 | 92% | 0.0100 | 0.0022 | 0.02 | 0.3151 | 34% |  | 64% | 69% |
|  | [13.9] | [39.9] | [45.2] | [2.9] | [31.7] | [5.0] | [7.6] | [23.9] | [24.7] |  |  |  |
| Formulation C 500 mg/kg  uninfected mice | 0.021 | 3.67 | 0.68 | 76% | 0.0048 | 0.0026 | 0.02 | 0.2555 | 27% |  | 49% | 79% |
|  | [23.9] | [0.5] | [49.6] | [5.4] | [75.9] | [30.9] | [12.6] | [42.8] | [83.5] |  |  |  |
| Formulation C 1000 mg/kg  uninfected mice | 0.011 | 0.18 | 0.51 | 99% | 0.0015 | 0.0085 | 0.03 | 0.1824 | 69% |  | 42% | 32% |
|  | [88.2] | [99.3] | [84.2] | [1.0] | [14.6] | [51.0] | [14.3] | [31.4] | [41.9] |  |  |  |
| Formulation A 125 mg/kg  Mtb-infected mice | 0.010 | 0.73 | 3.09 | 74% | 0.0028 | 0.0295 | 0.02 | 0.5223 | 11% |  | 139% | 92% |
|  | [38.5] | [116.9] | [0.5] | [35.9] | [13.8] | [244.5] | [11.6] | [31.0] | [479.5] |  |  |  |
| Formulation B 125 mg/kg  Mtb-infected mice | 0.007 | 0.91 | 3.09 | 83% | 0.0025 | 0.1620 | 0.01 | 0.4511 | 51% |  | 138% | 57% |
|  | [15.6] | [14.5] | [19.2] | [0.8] | [13.1] | [18.1] | [10.5] | [18.7] | [16.6] |  |  |  |
| Formulation B 250 mg/kg  Mtb-infected mice | 0.013 | 0.77 | 3.09 | 88% | 0.0025 | 0.0199 | 0.01 | 0.5338 | 11% |  | 131% | 91% |
|  | [70.9] | [62.7] | [6.9] | [7.8] | [11.4] | [81.9] | [12.1] | [40.2] | [683.1] |  |  |  |
| Formulation B 62.5 mg/kg  Mtb-infected mice | 0.005 | 0.63 | 1.70 | 77% | 0.0038 | 0.1906 | 0.01 | 0.0279 | 46% |  | 184% | 65% |
|  | [29.2] | [10.3] | [4.5] | [3.0] | [48.8] | [3.4] | [36.7] | [27.3] | [0.2] |  |  |  |
| Formulation C 125 mg/kg  Mtb-infected mice | 0.006 | 0.09 | 3.43 | 97% | 0.0212 | 0.0026 | 0.02 | 3.6279 | 54% |  | 128% | 48% |
|  | [25.2] | [44.4] | [0.0] | [1.2] | [25.3] | [5.0] | [10.3] | [0.5] | [20.7] |  |  |  |

**Table S5.** Scheme of the PK/PD study to evaluate the bactericidal activity of sorfequiline LAI formulations and oral control/comparator regimens, in the paucibacillary mouse model of TPT. For logistical reasons, the periods between immunization and infection; and infection and the start of treatment were truncated to 4 weeks instead of the standard ≥6 weeks. Study timepoints are relative to the start of treatment on Day 0. Blood samples (for measurement of sorfequiline and its M3 metabolite plasma concentrations) were obtained according to the sampling scheme presented in **Table S6**.

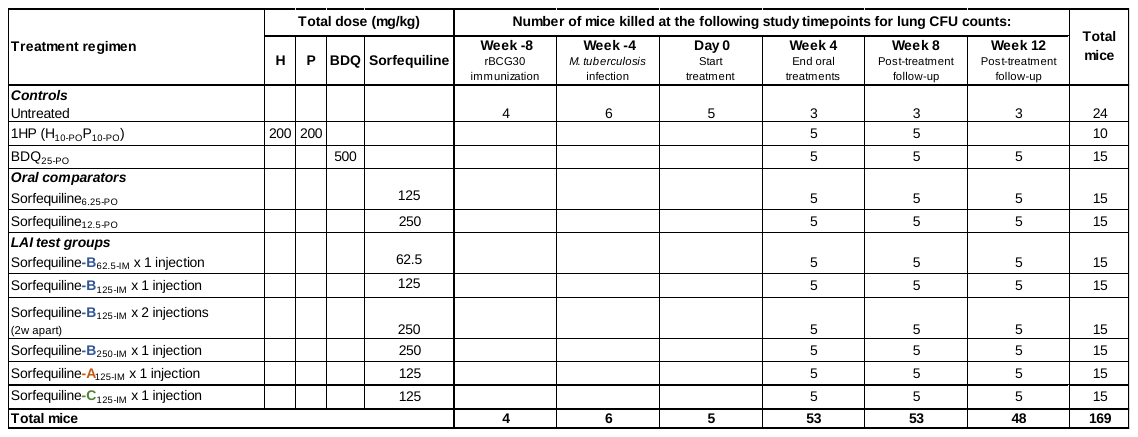

***Explanation of treatment regimen names***

*The subscript after each treatment agent indicates the dose in mg/kg followed by the method of administration: PO, oral gavage;	IM, intramuscular injection.*

*“1HP” indicates one month (i.e., 4 weeks) of oral isoniazid-rifapentine; this is a standard abbreviation for this regimen.*

*All oral regimens were administered 5 days per week (Monday to Friday) for 4 weeks.*

*For all LAI test groups, mice received an injection on Day 0; for mice that received 2 injections, the second injection was given after 2 weeks.*

**Table S6.** Scheme for blood sampling within the sorfequiline LAI PK/PD study for measurement of sorfequiline and its M3 metabolite concentrations in mouse plasma. Each treatment group consisting of a single injection of a sorfequiline LAI formulation included a total of 15 mice (M1 to M15). At each PK sampling timepoint, blood was obtained by in-life mandibular bleeding from sub-cohorts of three mice within each treatment group, and shading in the table is used to identify the 3-mouse cohorts. The complete study scheme with explanations of all treatment groups and total mouse numbers is presented in **Table S5**.

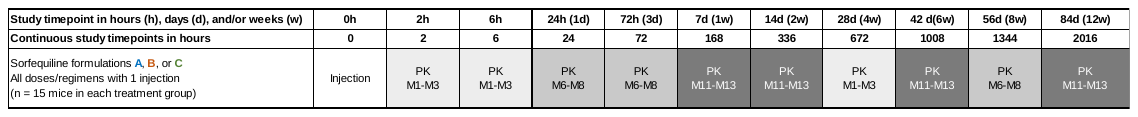

**Table S7.** Summary bactericidal activity (Mtb lung CFU) data for sorfequiline LAI formulations and control/comparator oral regimens in a mouse model of TPT. The complete study scheme with explanations of all treatment groups and total mouse numbers is presented in **Table S5**. Individual mouse CFU data are provided in **Supplementary Data** **S1D.**

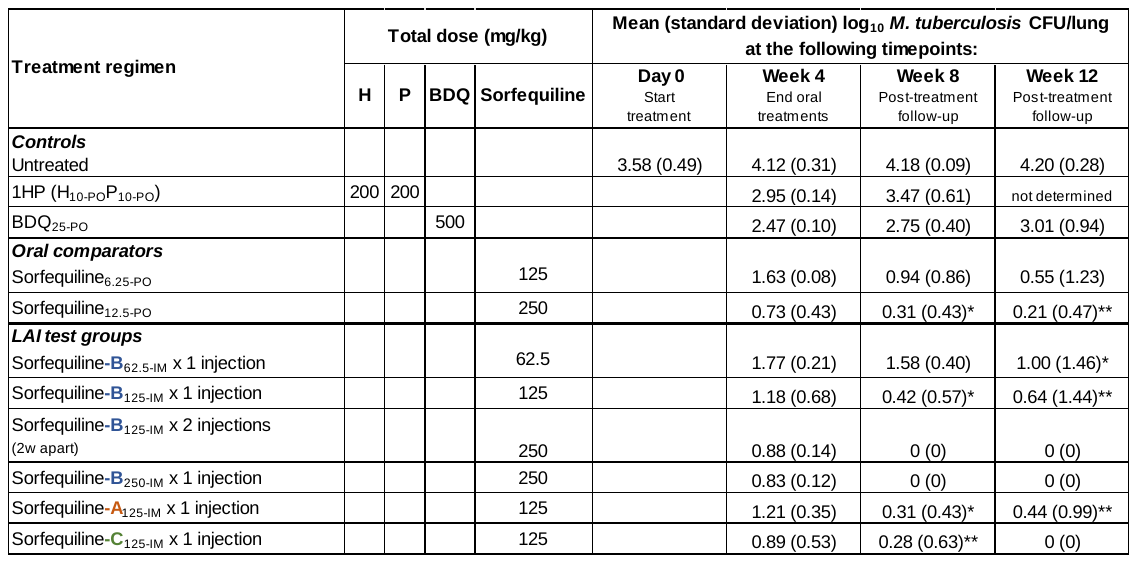

**Explanation of treatment regimen names**

- The subscript after each treatment agent indicates the dose in mg/kg followed by the method of administration: PO, oral gavage; IM, intramuscular injection.
- “1HP” indicates one month (i.e., 4 weeks) of oral isoniazid-rifapentine; this is a standard abbreviation for this regimen.
- All oral regimens were administered 5 days per week (Monday to Friday) for 4 weeks.
- For all LAI test groups, mice received an injection on Day 0; for mice that received 2 injections, the second injection was given after 2 weeks.

*Indicates that 3/5 mice were culture-negative

**Indicates that 4/5 mice were culture-negative

**Table S8.** Assessment of decreased susceptibility to DARQs in Week 12 lung homogenates from sorfequiline LAI PK/PD study. At Week 12, lung homogenates were also cultured on 7H11 agar plates containing 0.125 mg/L BDQ, and the ratio of CFU/lung cultured on BDQ-containing agar to the CFU/lung on plain agar was used as a proxy for the frequency of resistance to BDQ 0.125 mg/L (thus serving as a proxy for decreased susceptibility to DARQs). The plain agar contained activated charcoal, and the BDQ agar did not. Neither type of agar contained the selective drugs HYG or TCH, and thus CFU counts represent the total mycobacterial population (i.e., Mtb and rBCG30). Individual mouse data for Mtb and rBCG30 lung CFU counts are provided in **Supplementary Data** **S1D** and **S2B**, respectively. The complete study scheme with explanations of all treatment groups and total mouse numbers is presented in **Table S7**.

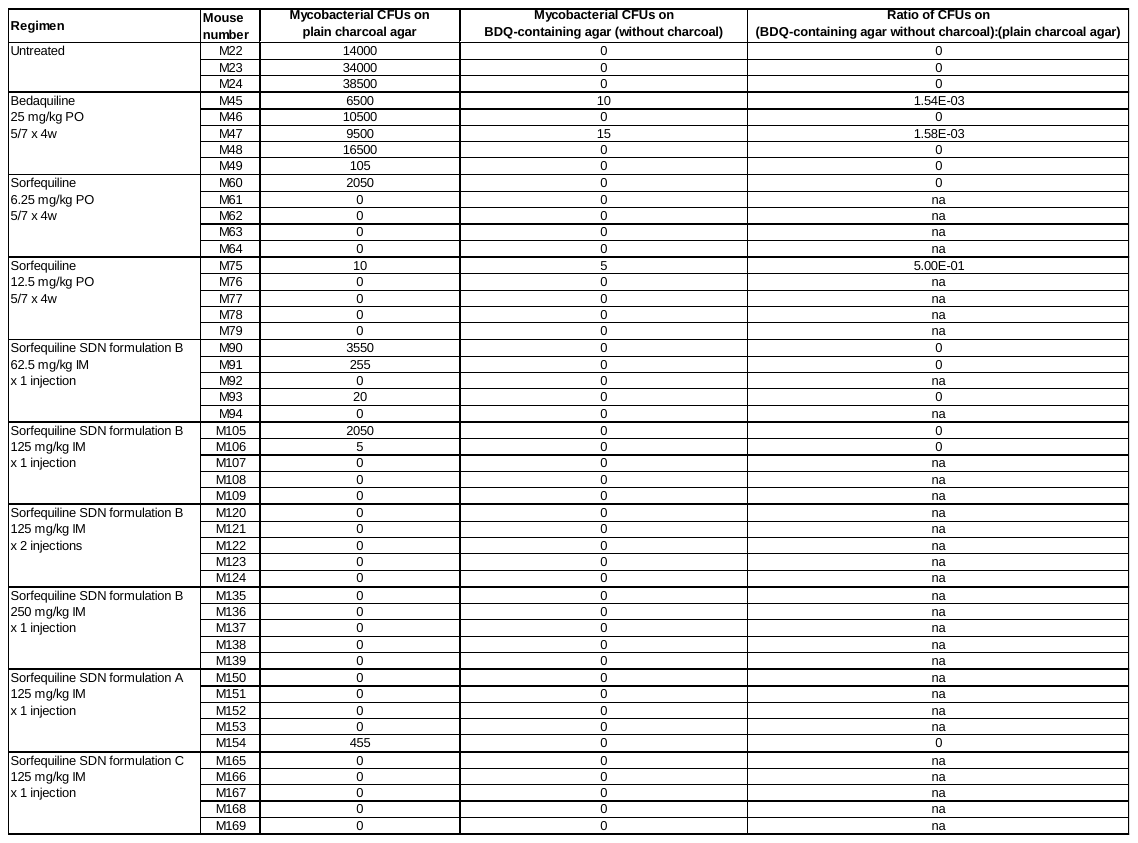

**Figure S1.** PK/PD data from study with oral dose-ranging of sorfequiline in a mouse model of TPT. **(A)** Lung CFU data for *M. tuberculosis* H37Rv at treatment initiation (Day 0) and after 4 weeks of treatment with the indicated treatment regimens. All drugs were administered by oral gavage, once daily, 5 days per week (Monday-Friday). 1HP = 1 month (4 weeks) of isoniazid 10 mg/kg and rifapentine 10 mg/kg. Bars and error bars represent mean and standard deviation, respectively (n = 4-6 mice per group). Dotted line indicates the mean CFU load in the untreated (comparator) group at Week 4. **(B)** Plasma concentrations of sorfequiline and its M3 metabolite after 23 days of treatment. Samples were collected just prior to (0 h) and 3 hours after the day’s oral dosing. Solid circles with solid lines indicate sorfequiline parent compound concentration, and open circles with dashed lines indicate M3 metabolite concentration. Data points and error bars represent mean and standard deviation, respectively (n = 3 samples per timepoint; the same mice were sampled at 0 h and 3 h). See experimental scheme in **Table S1**. All individual mouse data are available in **Supplementary** **Data S1A**.

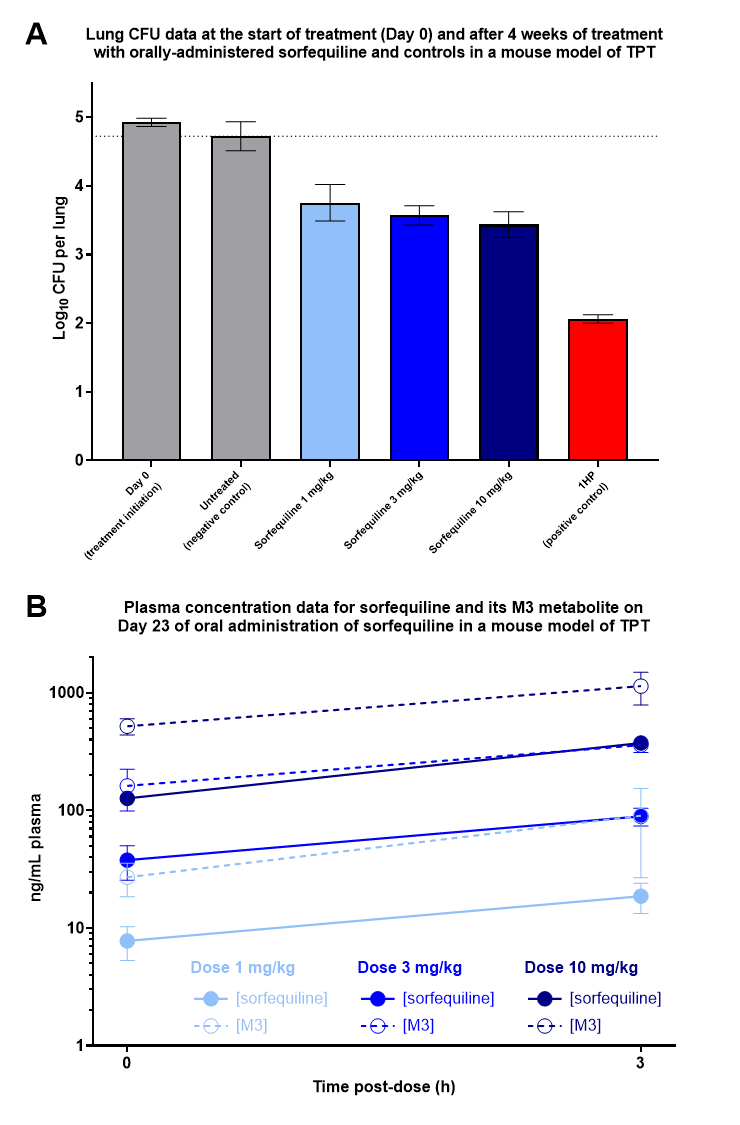

**Figure S2.** *M. tuberculosis* H37Rv lung CFU (PD) data from study with oral dose-ranging of DARQs bedaquiline, TBAJ-587, and sorfequiline in a mouse model of TPT. Lung CFU data are shown after 4 weeks **(A)** and 6 weeks **(B)** since starting treatment. D0 = Day 0, the time of treatment initiation; UT = untreated; 1HP = 1 month (4 weeks) of isoniazid 10 mg/kg and rifapentine 10 mg/kg. The numbers underneath the DARQ data bars indicate the daily dose in mg/kg. All drugs were administered by oral gavage, once daily, 5 days per week (Monday-Friday). Bars and error bars represent mean and standard deviation, respectively (n = 4-6 mice per group). Dotted line indicates the mean CFU load in the untreated (comparator) group at the same timepoint. See experimental scheme in **Table S2**. All individual mouse data are available in **Supplementary** **Data S1B**.

**A.	B.**

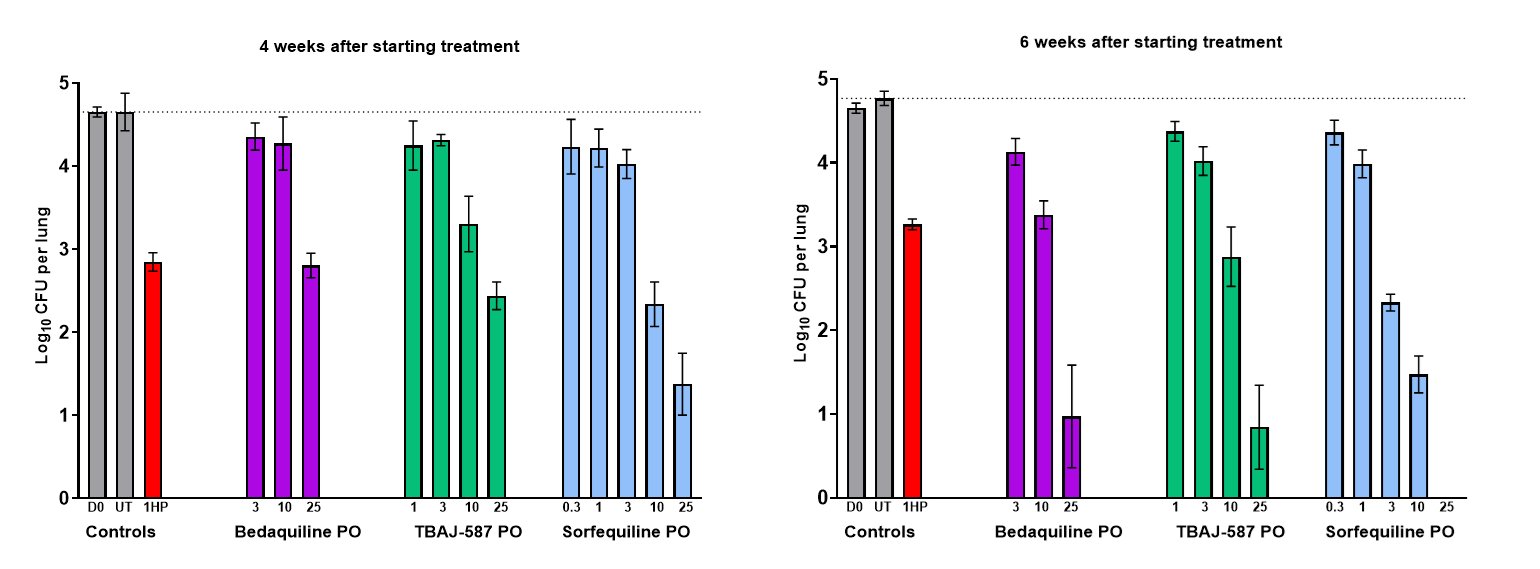

**Figure S3.** Schematic and differential equations for the compartmental PK model applied in simultaneous nonlinear regression fittings to observed plasma PK profiles for sorfequiline and its M3 metabolite. par: parent sorfequiline; met: M3 metabolite; Ax: amount; Cx: concentration; CLx: clearance; Vx: volume of distribution; Kx: first order rate constant; (643/657): Mw of M3 divided by Mw of sorfequiline = stoichiometric molar conversion of amount of parent to amount of metabolite; F2: fraction of dose assigned to slow release of sorfequiline; FAM fraction of F2 fraction assigned for depot release as M3 metabolite.

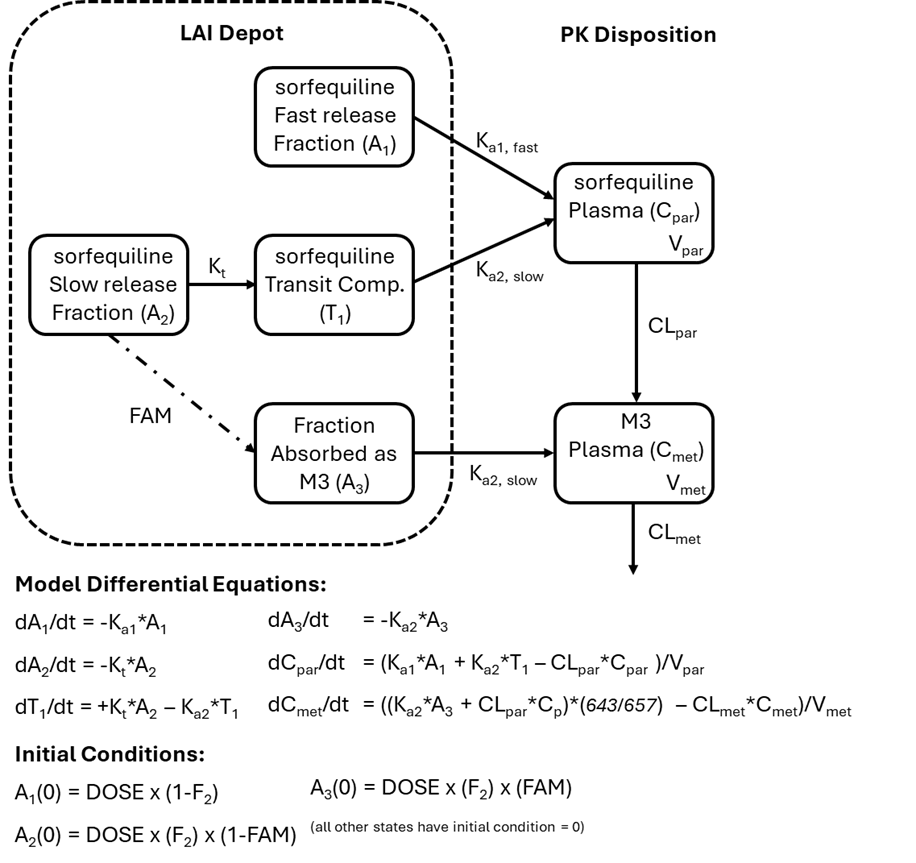

**Figure S4.** Plasma concentrations of sorfequiline (A-C), its M3 metabolite (D-F), and the combined total equivalent (eq.) of sorfequiline (G-I) following a single IM injection of sorfequiline LAI formulations A, B, and C, each at three dose levels in uninfected female BALB/c mice, raw data only. See scheme in **Table S3**. Individual mouse data are provided in **Supplementary Data S1C**.

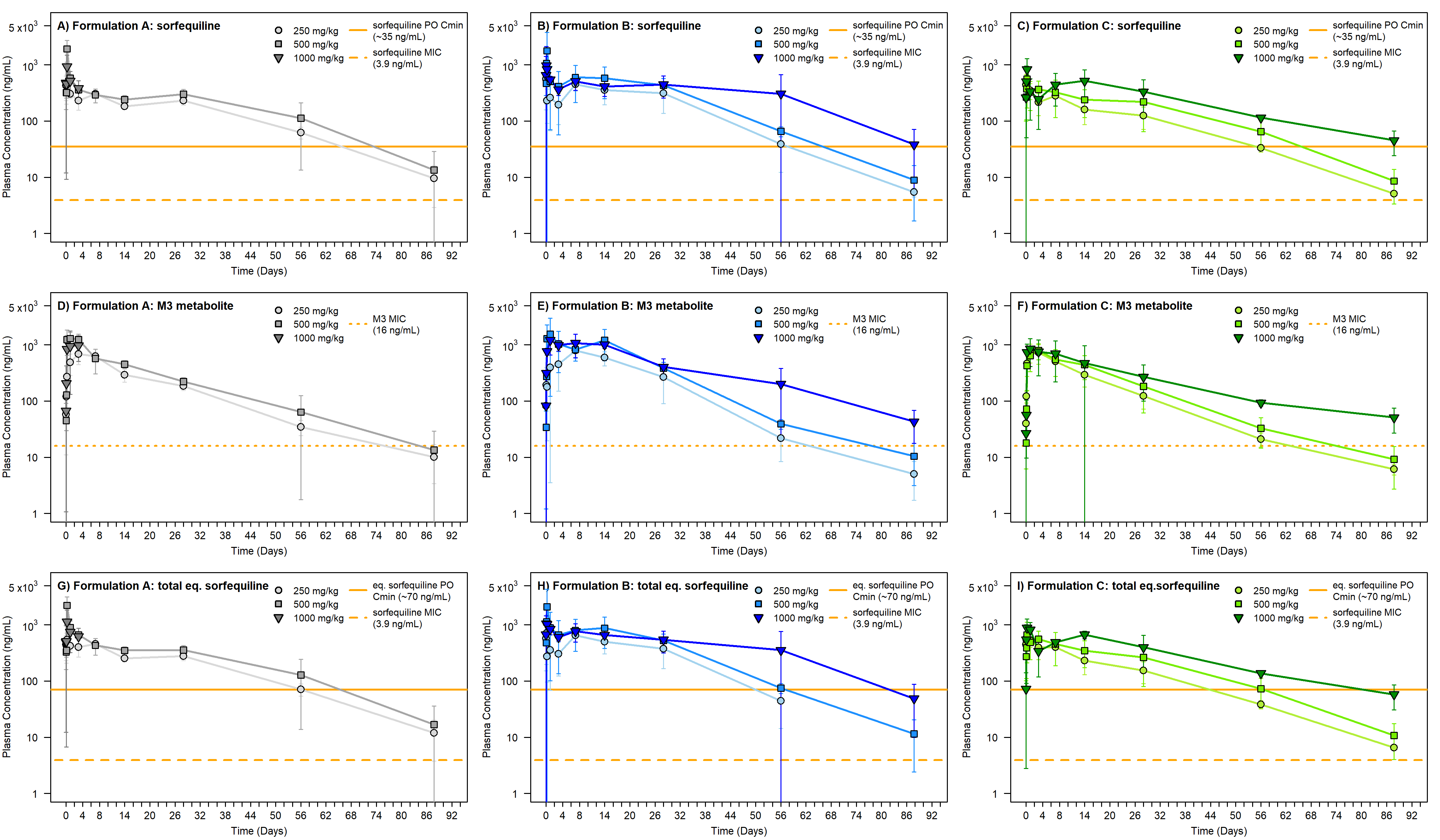

**Figure S5.** Lung CFU counts associated with treatment in the paucibacillary mouse model of TPT with a single injection of sorfequiline formulations A, B and C at 125 mg/kg (Panel A) and dose ranging of a single injection sorfequiline formulation B at 62.5, 125, and 250 mg/kg, and 2 injections at 125 mg/kg separated by 2 weeks (Panel B), with reference control oral treatment profiles. CFU data points represent mean values, and error bars represent standard deviation (n = 5 mice per timepoint). Matching associated plasma concentration profiles following these treatments (raw data only) presented for sorfequiline (panels C and D), its M3 metabolite (panels E and F), and the combined total equivalent (Eq.) of sorfequiline (panels G and H) in the TPT mouse model animals. PK data points represent mean values, and error bars represent standard deviation (n = 3 samples per timepoint). PK plots are illustrated with reference lines of provisional targets of oral C_min_ and MIC for sorfequiline parent, M3, or combined total equivalent of sorfequiline. Other abbreviations: Form, formulation, inj, injection. Study design and PK sampling schemes are provided in **Tables S5** and **S6**, respectively; individual mouse data are provided in **Supplementary Data S1D**.

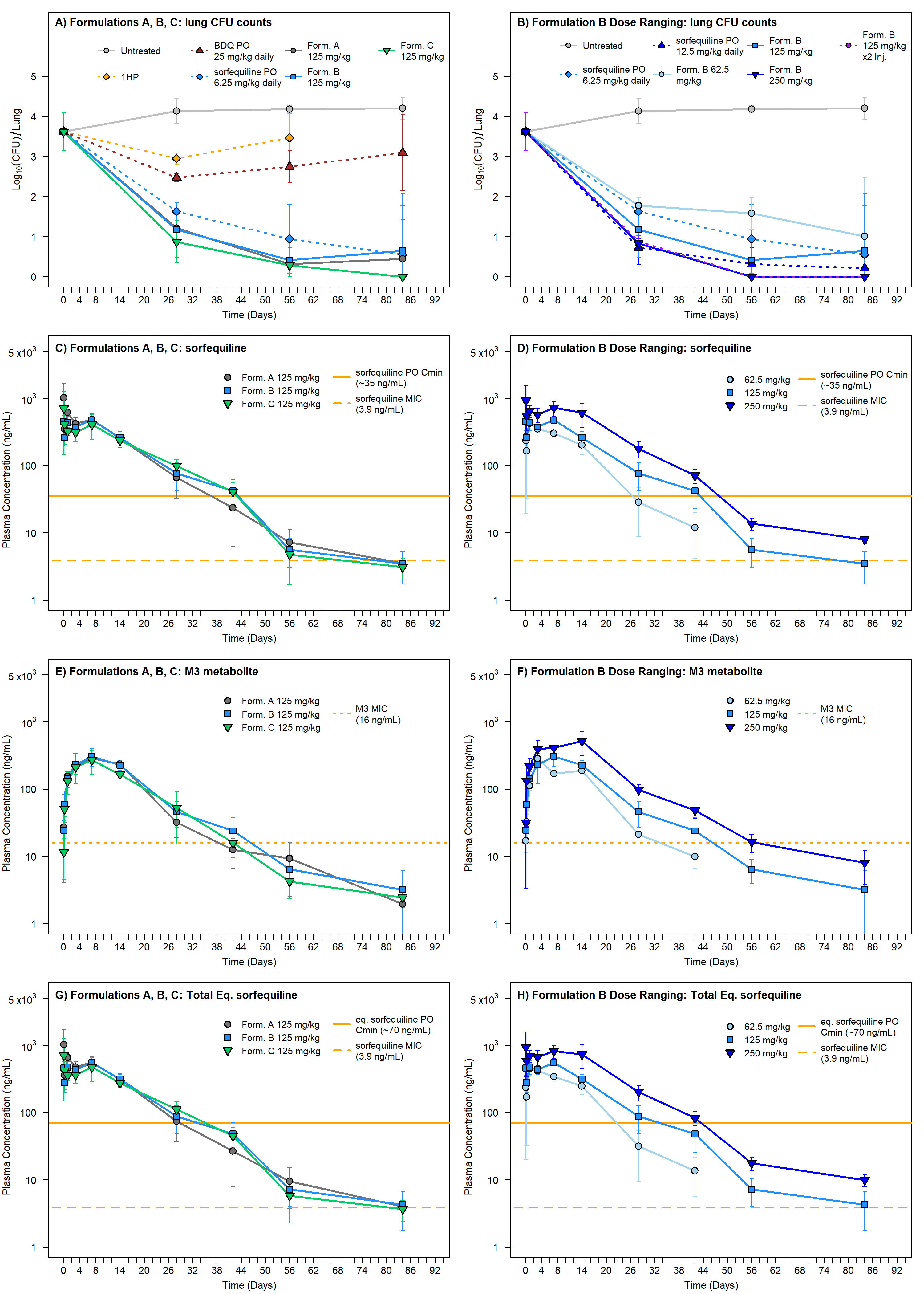
